# Molecular mechanism of claudin-15 strand flexibility

**DOI:** 10.1101/2021.12.07.471660

**Authors:** Shadi Fuladi, Sarah McGuinness, Le Shen, Christopher R. Weber, Fatemeh Khalili-Araghi

## Abstract

Claudins are one of the major components of tight junctions that play a key role in formation and maintaining epithelial barrier function. Tight junction strands are dynamic and capable of adapting their structure in response to large-scale tissue rearrangement and cellular movement. Here, we present molecular dynamics simulations of claudin-15 strands of up to 225 nm in length in two parallel lipid membranes and characterize their mechanical properties. The persistence length of claudin-15 strands is comparable with experiments leading to a curvature of 0.12 nm^−1^ at room temperature. Our results indicate that lateral flexibility of claudin strands is due to an interplay of three sets of interfacial interaction networks between four linear claudin strands in the membranes. In this model, claudins are assembled into interlocking tetrameric ion channels along the strand that slide with respect to each other as the strands curve over sub-micrometer length scales. These results suggest a novel molecular mechanism underlying claudin-15 strand flexibility. It also sheds light on the inter-molecular interactions and their role in maintaining epithelial barrier function.

## Introduction

Tight junctions are belt-like, circumferential cell-to-cell contacts between epithelial cells that regulate the passage of water, ions, and small molecules through paracellular space^1–5^. Tight junction proteins polymerize to form a two dimensional network of branching strands in each cell membrane and pair with tight junction strands in adjacent cells^5–9^. Tight junction transmembrane proteins include tetra-spanning proteins such as claudins and occludin, and other mono and tri-transmembrane domain containing proteins^19–22^.

Claudins are one of the major components of tight junctions that play a key role in tight junction assembly and are responsible for selective paracellular permeability^2,5,10,23–29^. Based on studies where claudins are expressed in fibroblasts it has been shown that claudin strands are dynamic with structural flexibility to bend, arch, and to form new branches on the scale of nanometers to micrometers^10–12^. The structural flexibility of claudin strands is important in maintenance of barrier function during changes in cellular morphology and tissue rearrangement^13–15^. In fact, variations in strand morphology can influence tight junction barrier function^7,16–18^.

Claudins are transmembrane proteins with four transmembrane helices (TM1-TM4) that form a left-handed bundle and two extracellular segments (ECS1 and ECS2). The extracellular segments consist of an extracellular helix (ECH), five antiparallel *β*-strands, and a highly conserved consensus W-LW-C-C motif^27,30–32^. Both ECS1 and ECS2 are essential for claudin assembly and charge selectivity of the pores^2,32–39^.

The crystallographic structure of claudin-15 shows that claudin-15 forms a linear polymer^32^. The authors also suggest that the structurally disordered loops in ECS1 and ECS2 are vital for tight junction pore formation. However, the structure of these loops was not revealed. Based on cysteine cross-linking experiments, Suzuki et^34^ al proposed a set of inter-molecular *cis-*interactions mediated by the *β*-sheets of ECS1 (here termed “face-to-face”) that lead to a continuous arrangement of claudins into half-pipe structures^34^. Side-by-side *cis-*interactions between claudins in the same membrane form the strand backbone. The paracellular pore structure is then completed through head-to-head *trans-*interactions between extracellular loops of claudins on adjoining cells. This model was verified in atomic simulations confirming the head-to-head interactions between claudin loop domains in the adjoining cells and elaborating on the molecular basis for paracellular ion channel selectivity^40–43^. However, the proposed model by Suzuki et al.^34^, does not account for the different morphologies of tight junction strands observed under freeze-fracture electron microscopy including strands with curved structures at the micrometer length scales, and cannot explain the protein-protein interactions necessary for strand flexibility. Longer strand models and dynamics studies are needed to identify the molecular basis for claudin-15 strand flexibility.

In this study, we used a refined atomic model of claudin-15 paracellular ion channels^40^ to simulate claudin strands at scales previously unexplored. We carried out molecular dynamics (MD) simulations of claudin strands ranging from 30 to 225 nm in length, with 36 to 300 monomers, in two parallel lipid membranes. We investigated their mechanical properties at microsecond timescales. To achieve longer time scales, we simulated the strands at all-atom as well as hybrid resolution, where the protein is represented atomically and lipid and water molecules are coarse-grained^44–46^. We successfully visualized the curvature of claudin-15 strands, calculated their persistence length, and identified intermolecular interactions that lead to their flexibility. These findings suggest a novel mechanism for bending flexibility of claudin strands that is consistent with experimentally observed curvatures of tight junctions strands.

## Results and discussion

We used MD simulations to study the stability and mechanical properties of claudin strands consisting of claudin-15 pores in two parallel lipid membranes. We constructed atomic models of claudin strands with lengths ranging from 30 nm to 225 nm consisting of 36 to 300 claudin-15 monomers. These models are based on our previously refined model of three claudin-15 pores with twelve monomers^40^, which was replicated to create longer claudin strands. The functional characteristics of claudin-15 pores in this model were previously verified, showing that it recapitulates experimentally defined size and charge-selectivity of claudin-15 channels^40^.

Our molecular dynamics modeling succesffully modeled ion transport through the claudin-15 pore, but it was not able to model the long term interactions and dynamics among claudin-15 monomers. To address this questions,, we simulated the systems using a hybrid-resolution model, PACE, that combines a united atom representation of proteins with a coarse-grained model of lipids and solvent (MARTINI) to achieve longer time scales and better sampling of the configuration space, which is necessary to study strand organization in the membran^44–46^. To validate the results of the PACE forcefield, structural stability of claudin monomers, their fluctuations, and the integrity of claudin pores in the 36-claudin strands (30 nm long) are compared against a 250 ns all-atom trajectory of the same system. We then characterized mechanical properties of the strands as a function of their length and determined inter-molecular contacts between claudin monomers that confer flexibility to the strand.

### Stability of claudin pores

A model of 36 claudins in two parallel POPC lipid membranes was constructed (Fig. 1a) based on our previously refined three-pore model of claudin-15. In this model, each lipid bilayer contains an antiparallel double-row of claudin monomers arranged linearly through two sets of side-by-side (*cis-*) and face-to-face interactions (Fig. 1b). Head-to-head (*trans*-) interactions with claudins in the second lipid membrane then result in formation of claudin channels (pores) in the space between the two membranes. Each pore in this model is made of the assembly of four claudins, two from claudins in each membrane.

**Figure 1:**
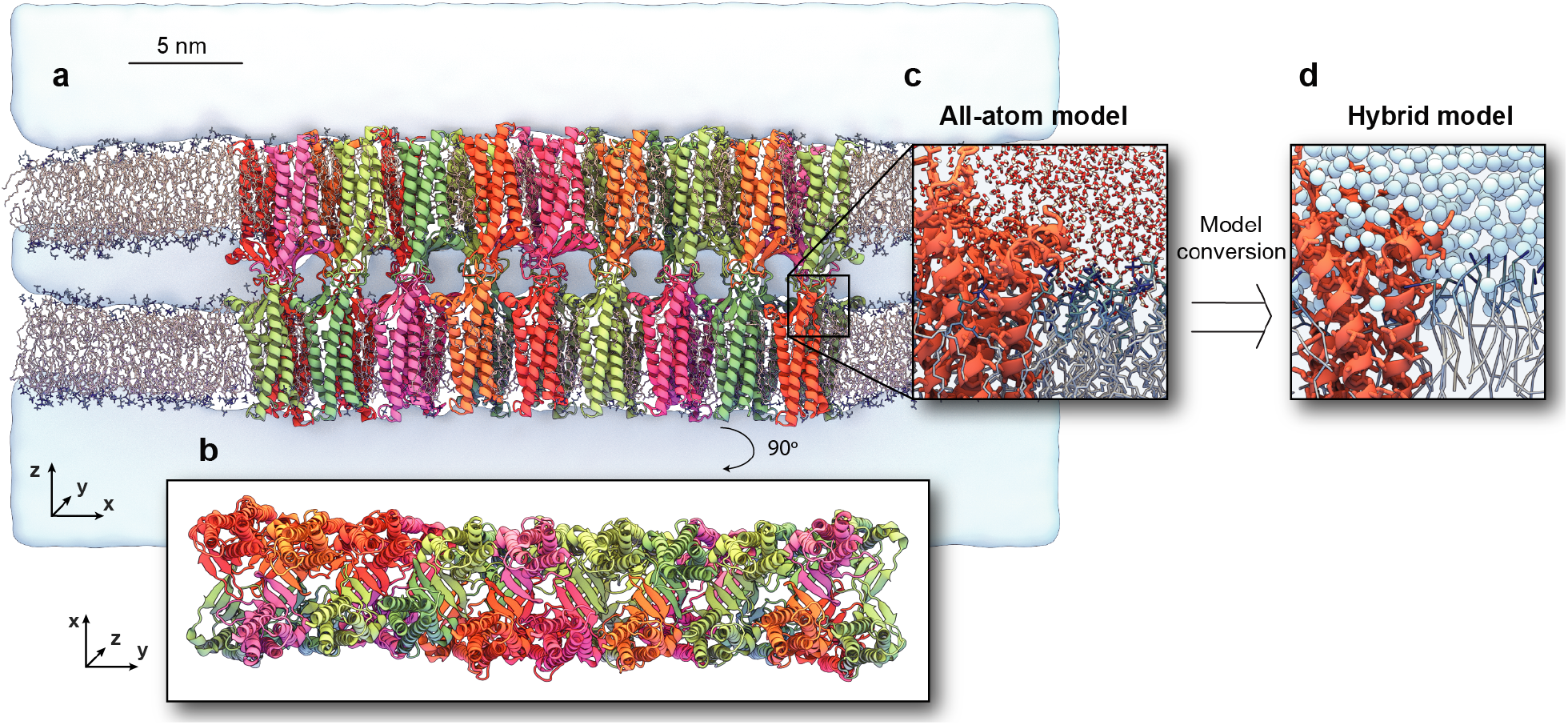
Claudin-15 strand structure. (a) All-atom model of a claudin-15 strand containing 36 claudin monomers embedded in two parallel POPC bilayers and surrounded with solvent. (b) Top view of the system shows claudin monomers association in a double row arrangement. (c) This simulation system is prepared with using an atomic-resolution forcefield. (d) A hybrid-resolution model of the system is created by converting the lipid and solvent molecules to a coarse-grained representation using the PACE forcefield. The hybrid model preserves the atomic structure of the protein, while models lipid and solvent molecules with a coarse-grained representation.

The system of 36 claudins was equilibrated for 250 ns with an all-atom representation of the system including proteins, lipids, water, and ions (Fig. 1c). The system remained stable with an average root-mean-square deviation of 2.19 ± 0.49 Å with respect to the initial structure of the backbone (Fig. 2a). Face-to-face interactions between claudins in each membrane were maintained through hydrogen-bonds between *β*-sheets (ECS1) of neighboring claudins and were present for ∼63 % of the time. The stability of the system was comparable to the smaller systems of claudin strands^40,41^.

**Figure 2:**
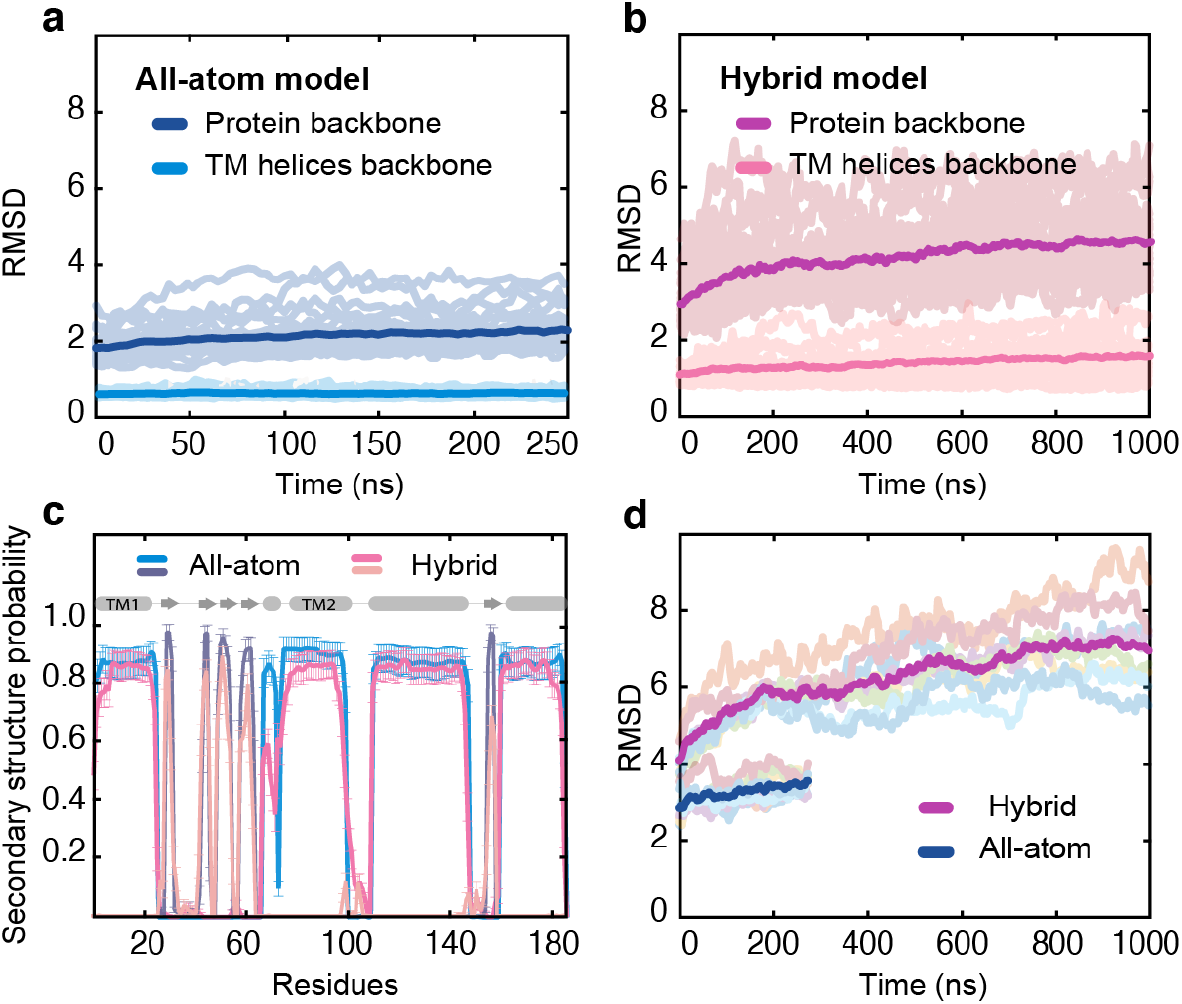
Structural stability of claudin-15 strands: Root-mean-square deviation (RMSD) values of the protein backbone and transmembrane (TM) helices backbone with respect to the initial structure throughout the simulation trajectory of (a) all-atom and (b) hybrid model. Average of RMSD values of the 36 monomers are shown in bold lines. (c) Secondary structure content of claudin-15 monomers, averaged over 36 monomers and the trajectory time in all-atom (250 ns) and hybrid (1 *µ*s) models of claudin-15 strands. (d) RMSD values for the backbone of tetrameric claudin channels with respect to the initial structure and the average over eight channels.

In addition to this all-atom simulation, the 36-claudin system was simulated in a hybrid environment where the lipids and solvent molecules were coarse-grained and the protein was represented atomically^44–46^ (Fig. 1d). During this 1 *µ*s-long simulation, the claudin strand remained stable with an average root-mean-square deviation of 4.21 ± 0.88 Å for protein backbone with respect to the initial structure (Fig. 2b). Further comparison of the two simulation trajectories indicates that the secondary structure of claudin monomers is well-preserved in the hybrid-resolution model (Fig. 2c). The tetrameric structure of claudin pores is also maintained in both simulations, with average backbone root-mean-square deviations of 3.86 ± 0.30 Å and 6.56 ± 0.89 Å for all-atom and hybrid resolution trajectories after fitting the pores to the initial structure (Fig. 2d).

### Claudins form water channels

Claudins form tetrameric ion channels that run parallel to the membrane surface and seal the paracellular space (between two membranes) to water and ions. Simulations of the 36-claudin system show fully hydrated channels formed by extracellular domains of claudins that are ∼40-50 Å long (Fig. 3a). The average linear density of water molecules along the pores is 7.5 water molecules per Å corresponding to an effective radius of 7.2–9.2 Å. The difference between the effective radii of the pore in all-atom and hybrid-resolution models is due to the coarse-grained nature of water molecules in the latter, where four water molecules are represented as one molecule^47^.

**Figure 3:**
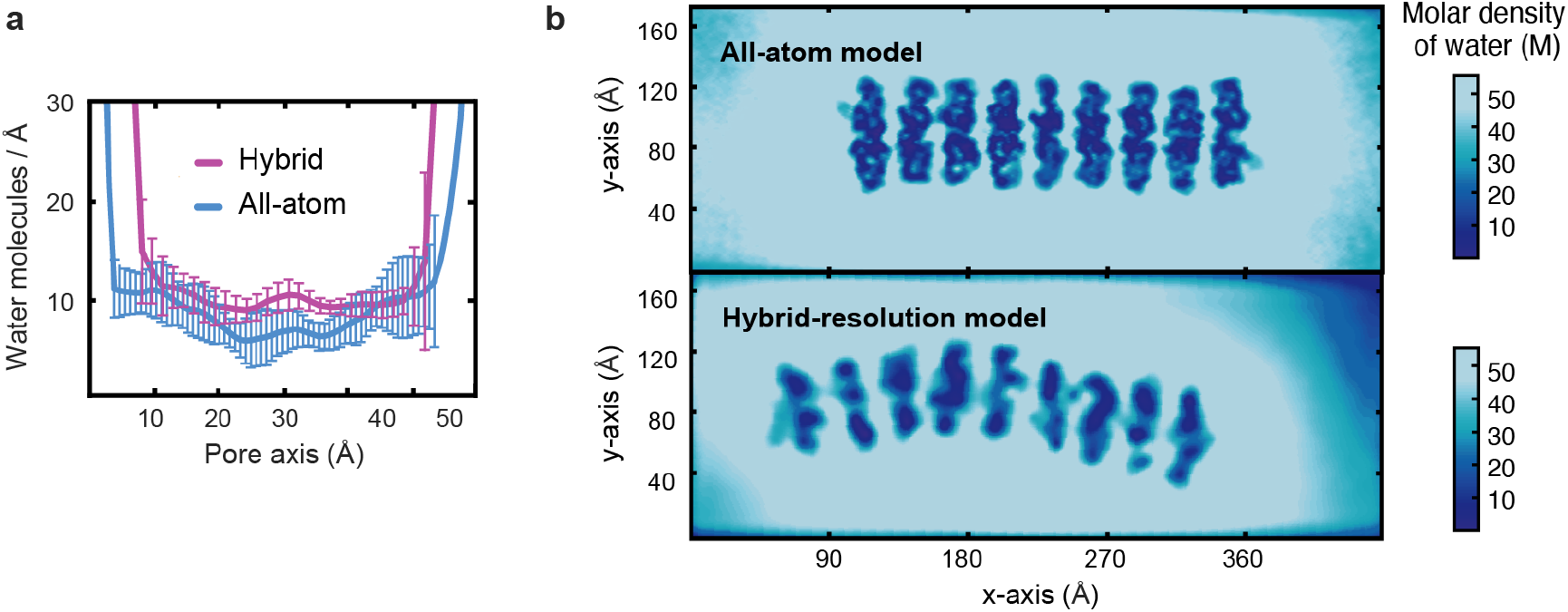
Water density across paracellular space shows integrity of channel structure: (a) Number of water molecules along the length of channel, averaged over eight channels in each strand (of 36 monomers and 300 Å -long), and over 30 ns of equilibration simulation. Number of water molecules in hybrid model is multiplied by four to account for coarse-grained water mapping. (b) The averaged water density from simulation trajectory (30 ns) across a plane parallel to the membranes and crossing the channels in the middle, for all-atom and hybrid resolution models.

Figure 3b shows the density profile of water molecules on a cross-sectional plane in the middle of two membranes. Eight water-filled channels run parallel to this plane, separated by regions with no water presence (dark blue). The channels are filled with water molecules at a density close to the bulk and are separated by extracellular domains (ECS1 and ECS2) of claudin. Head-to-head (*trans-*) interactions between ECS1 and ECS2 of claudins in two opposing membranes seal the space between the two membranes and limit the flow of water and ions to claudin pores. Non-specific hydrophobic patches corresponding to the *variable* regions on ECS1 and ECS2 are responsible for head-to-head interactions of claudins^24,36,37,48–50^. These interactions are well-maintained during the simulations and form well-defined pathways for ions and water in paracellular space despite the dynamic nature of the strands. Dynamic curvature of the strands is more apparent in the hybrid-resolution trajectories, where claudin tetramers forming ion channels adapt to the local curvature of the strand and maintain the pore integrity through persistent head-to-head and face-to-face interactions.

### Mechanical properties of claudin-15 strands

To characterize mechanical properties of claudin strands, three additional systems of 84, 180, and 300 claudins corresponding to strands of 63 nm, 135 nm, and 225 nm in length were prepared. Each system consists of two double-rows of claudins in two parallel POPC lipid membranes forming 20, 44, or 74 ion channels (Fig. 4a). Each system was simulated for 1 *µ*s in the hybrid-resolution representation, and the stability of the system was assessed similar to the smaller system (summarized in Table SI and SII in the supplementary material). The persistence length and bending modulus of claudin strands in the plane of membrane were calculated from thermal fluctuations of the system at 300 K.

**Figure 4:**
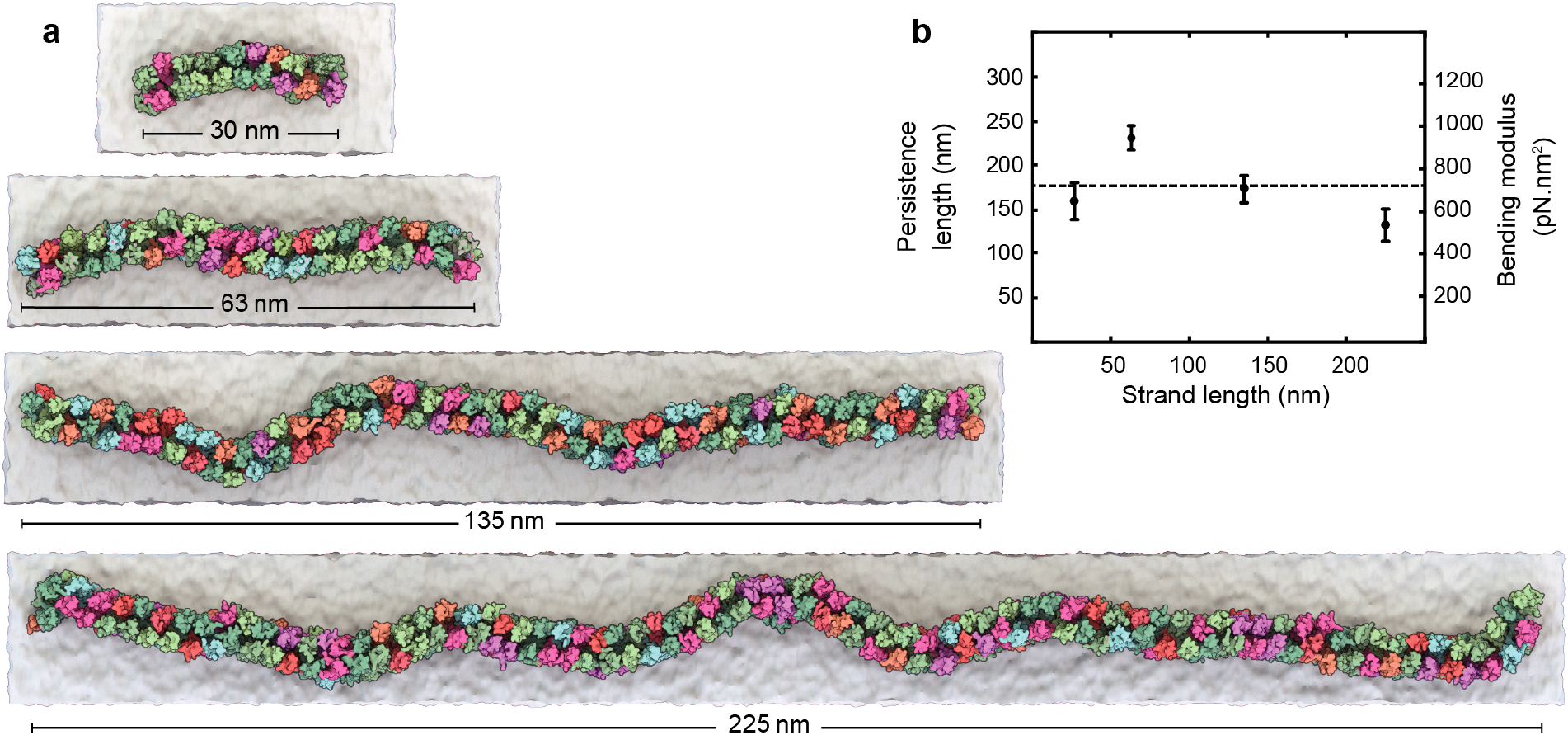
Claudin-15 strands adopt a “curvy” morphology: (a) Equilibrated configuration of strands of various lengths show the “curvy” morphology of claudin-15 strands despite the “straight” arrangement of initial model. (b) The persistence length (left-hand axis) and the bending modulus (right-hand axis) are measured for each length of claudin-15 strands. The horizontal line shows the average of measurements.

The persistence length of the strands *l*_*p*_ is an indication of the stiffness (or flexibility) of the strand in the plane of membrane and is defined as the length over which the direction of the strand becomes uncorrelated. The persistence length of claudin strands is calculated from equilibrium simulation trajectories. Assuming that the correlation function between tangent vectors at two points on the strand decays exponentially as a function of their separation, the persistence length is calculated as the characteristic length scale of this exponential decay along the strand. The persistence length of claudin-15 strands was estimated from equilibrium trajectories of four strands and corresponds to an average of *l*_*p*_ = 174 nm with a standard deviation of 42 nm among four systems (Fig. 4b). This is consistent with persistence length of 191 ± 184 nm estimated from freeze-fracture electron microscopy images of claudin-15 strands^12^.

Persistence length of the strands *l*_*p*_ is a function of temperature and bending rigidity of the strands, which is usually described by its flexural force constant *κ*_*b*_. The bending rigidity determines how much energy *E*_*b*_ is stored in a strand of length L upon bending to an angle of *θ*, and is a function of strand geometry as well as its Young modulus:

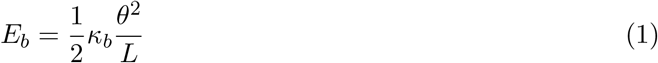

Thermal fluctuations in the local bending angles of the stand are then related to the bending rigidity of the stands, which in turn defines the persistence length *l*_*p*_ of the strands at each temperature T as:

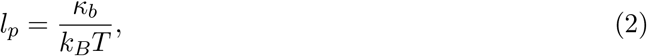

where *k*_*B*_ is the Boltzmann constant.

Bases on the persistence length *l*_*p*_ of the claudin-15 strands obtained from equilibrium trajectories, we estimated the bending rigidity *κ*_*b*_ of the strands as 715 ±174 pN.nm^2^ (Fig. 4b). To put this value into context, approximately, 0.24 *k*_*B*_T energy is needed in order to bend a 10 *µ*m long claudin-15 strand through an angle of 30° at 300 K. This bending rigidity is comparable to the rigidity of double-stranded DNA with a persistence length of 50 nm^51–56^, is four orders of magnitude smaller than the bending rigidity of force-transmitting microtubules^57–61^, and one order of magnitude smaller than that of actin filaments with persistence lengths in mm and *µ*m range^57,62,63^. This is consistent with dynamics nature of claudin strands as seen in live florescence imaging^10,12,64^.

Coarse-grained (CG) and all-atom simulations are commonly used to calculate mechanical properties of biological systems, e.g., lipid membranes, from molecular dynamics trajectories. For example, the bending modulus of POPC lipid bilayers is reported as 29-32 *k*_*B*_T from all-atom simulations based on the CHARMM36 forcefield^65,66^ and 20-25 *k*_*B*_T from coarse-grained simulations using the MARTINI forcefield^66,67^. Both forcefields consistently reproduced experimentally reported values of the POPC bending moduli which are reported to be in the range of 5.8-49 *k*_*B*_*T*^68,69^. Hence, it is expected that the hybrid resolution model used here reasonably produces the persistence length and bending rigidity of claudin-15 strands.

### Flexibility mechanism of claudin-15 strands

The bending flexibility of claudin strands in the membrane stems from flexible arrangement of monomers within the membrane. Upon bending, adjacent claudins in each row deviate from their initial arrangement^32,34^ and are displaced to adapt to the local curvature. To characterize the dynamic behavior of claudin strands, we looked at the relative orientation of claudin pairs as the local curvature of the strands changes. We calculated the relative orientation angle of claudins as they rotate with respect to each other to accommodate local changes (side-by-side interactions) (Fig. 5a-c) as well as the distance between opposing claudin pairs in the double-row arrangement of claudins in each membrane (face-to-face interactions) (Fig. 5d-e).

**Figure 5:**
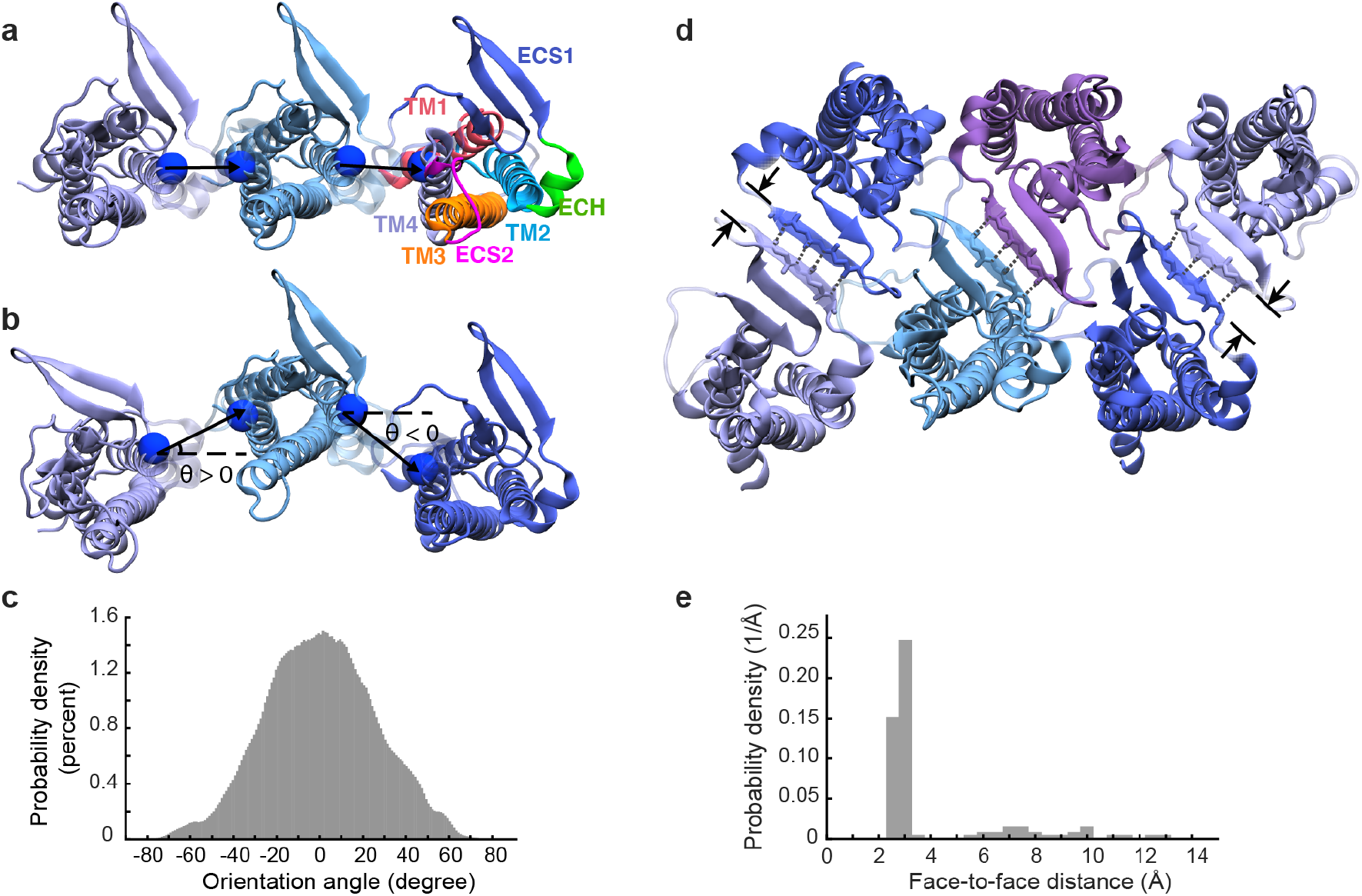
Relative orientation of claudin-15 monomers in strand simulations. The relative orientation of adjacent claudin monomers in (a) the straight (from x-ray crystallography) and (b) the curved arrangement of claudin-15 strands. The relative orientation of neighboring claudins is calculated as the tangent angle of the vector connecting the mid-points of the TM2 and TM4 helices of the two adjacent monomers relative to the initial arrangement. (c) The probability distribution of relative orientation angles of adjacent claudins in the 225 nm-long (300 monomers) strand over the last 600 ns of simulation time. (d) The double row arrangement of claudin strands are stabilized through hydrogen bonds between the backbone of the fourth *β* strand in the extracellular segment (ECS1) of two claudins. The nitrogen and oxygen of the backbone of residues 60 to 64 that form a hydrogen bond are shown in blue and red. (e) The probability distribution of face-to-face distance of claudins is calculated as the minimum distance between the blue and red spheres (shown in (d)) in the 225 nm-long (300 monomers) strand over the last 600 ns of simulation time.

The distribution of relative orientation angle between claudin pairs in the 225 nm strand (300 claudins) shows a symmetric distribution of angles between –70° and +70° (Fig. 5c). The peak at the center at 0° corresponds to the linear arrangement of claudins observed in the crystal structure of claudin-15^32^. At the temperature of 300 K, 95% of adjacent monomers adopt a relative orientation of –50° to 50° and 68% have a relative angle of –26° to 26°. The width of the orientation distribution indicates that curvatures up to 0.29 nm^−1^ are permissible in claudin-15 double strands (based on the average distance of 30 Å between centers of mass of adjacent claudins), comparable with the analysis of claudin-15 strands freeze-fracture images^12^. This data suggests that the side-by-side (*cis-*) interactions, present in the initial model, are highly flexible and support the range of curvatures observed in this work.

We further characterized the atomic interactions between claudin pairs forming side-by-side (*cis-*) interactions from the simulation trajectories. Based on the relative orientation angle of claudins *θ* (Fig. 5a-c), we grouped claudin pairs into three clusters; a cluster corresponding to the linear arrangement observed in the crystal structure (–10° < *θ* < +10°), another cluster corresponding to the dominant positive orientation angles (+10° < *θ* < +30°) and the last cluster corresponding to the dominant negative orientation angles (–30° < *θ* < –10°). The three clusters represent three dominant interfaces between neighboring claudin pairs. Contact maps between claudin pairs show the main residues involved in the side-by-side interaction of claudins in each cluster (Fig. 6a-c). At each of these clusters three sets of interactions/regions are identified (colored red, purple, and orange in Fig. 6). These sets include interactions of the extracellular helix ECH; residues P66, S67, M68, L69, and L71, with three distinct regions: (I) ECS2/TM4; residues K155, E157, L158, and Y163 (ECH–ECS2; purple cluster), or (II) TM3; residues T143, F146, and F147 (ECH–TM3; orange cluster), or (III) ECS1; residues T33, V34, and H35 (ECH–ECS1; red cluster). The probability of observing each of these interactions varies over time as the orientation angle between claudins changes.

**Figure 6:**
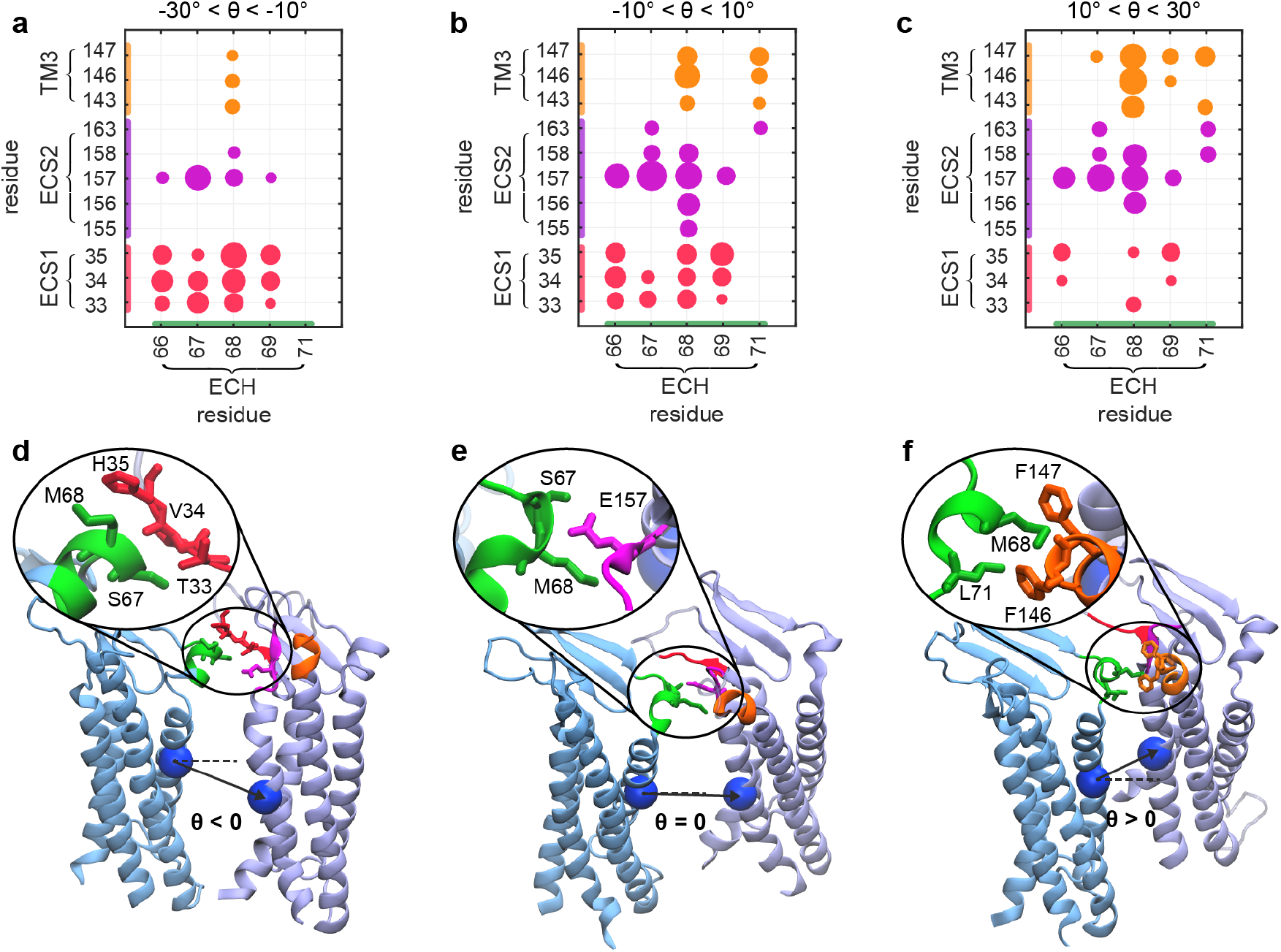
Multiple side-by-side interaction interfaces provide the flexibility of claudin-15 strands. (a-c) Contact maps of the side-by-side interactions between claudin pairs grouped into three clusters based on the pairs relative orientations. The size of the circles denotes the probability of the pair-wise interactions and the colors correspond to the interacting region. The analysis is performed over the 225 nm strand (300 monomers) and 15 ns. The snapshots of dominant interactions at each cluster are shown for claudin pairs corresponding to (d) negative, (e) zero, and (f) positive orientation angles. The interacting regions are colored by green for ECH, red for ECS1, magenta for ECS2, and orange for TM3.

The most dominant interaction is between residues S67/M68 of ECH and E157 of ECS2 (region I) consistent with previous reports^12,40^. E157 is a highly conserved residues in claudin family^35–37,49^ and in this model is located at the entrance of the pore^40^. Mutation of E157 is shown to disrupt strand formation^12^. A second dominant interaction is between M68 of ECH and F146/F147 of TM3 (region II), which is observed mostly at the zero or positive orientation angles (Fig. 6b-c).

These interactions are reported in the crystal structure of claudin-15^32^ as the main connection between neighboring claudins for maintaining the *cis-*interactions. Mutation of M68, F146, and F147 to smaller or charged residues is shown to disrupt strand formation^32^. We also observed a third dominant interface between ECH and ECS1 in the simulation, which involves the variable region of ECS1 residues 33–35 (region III). This region was not resolved in the crystal structure of claudin-15, and was modeled in our claudin-15 pore^40^. This set of interaction is mostly observed at zero or negative orientation angles (Fig. 6a-c). The *β*1–*β*2 loop on ECS1 (32–42) is hypothesized to form strong hydrophobic seal in *trans-*interactions^31,32,70^ and MD simulations have identified strong pair-wise hydrogen bonds between residues 39–42 (the other end of the loop) of opposing claudins^40^. Therefore, residues 33 to 35 are accessible for *cis-*interaction with the adjacent claudins. Snapshots of the three interfaces and the residues involved are shown in Fig. 6d-f.

These findings suggest that the extracellular helix ECH acts as a pivot point around which claudin monomers rearrange and rotate around a centrally positioned ECH–ECS2 interface with S67–E157 as the dominant interaction site. At positive curvatures, corresponding to positive orientation angles (Fig. 6c) the frequency of contacts between ECH and ECS1 decreases and more contacts are established between ECH and TM3. Similarly, as the local curvature of strands changes and relative orientation angle between neighbors becomes negative, the interactions between ECH and ECS1 become more dominant (Fig. 6a). Spatial arrangement of ECS1 and TM3 on the two sides of ECS2 allows the strands to bend while maintaining side-by-side contacts. These three interfaces, providing a wide range of permissible orientation angles for side-by-side interaction, are critical in conferring flexibility to claudin strands.

In addition to the side-by-side interactions between claudins, we characterized the face-to-face interactions between claudins monomers in the membrane (Fig. 5d). The double-row arrangement of claudins involves an anti-parallel double row of claudins in each membrane that is stabilized through a network of four to six hydrogen bonds between *β*-sheets of claudins (face-to-face interactions)^40^. These hydrogen bonds are responsible for maintaining the tetrameric pore structure and for forming a well-defined permeation pathway. We calculated the average distance between the two *β*-strands of claudins over the simulation trajectories of the most dynamic strand with 300 monomers. The average distance between the two monomers over the last 15 ns of simulation was 3.84 ± 2.38 Å, indicating that face-to-face interactions are well-maintained (Fig. 5e). This data indicates that the two claudin rows in each membrane move together and explains why the local curvature of the two rows is correlated in the simulation trajectories.

These findings are recapitulated in a schematic representation (Fig. 7) showing the two dimensional movement of claudin monomers in the membrane with respect to each other. The arrows indicate the directional movement of tetrameric claudin channels (only a pair is shown here for clarity) in the plane of membrane. Claudin pairs, coupled to each other through face-to-face interactions, slide with respect to each other in the plane of membrane to produce a local curvature in the strands. Flexible side-by-side interactions between claudins is key in this sliding motion. Three sets of side-by-side interactions between ECH and ECS1, ECS2, or TM3 facilitate this movement. ECH shown in green acts as a pivot point in maintaining the side-by-side interactions, while claudins slide in the membrane with respect to each other. During this movement, the tetrameric structure of claudin ion channels is maintained by head-to-head interaction of the claudins in this strand with those in a second strand on top (not shown here for clarity). Simulations show that the local curvature of the strands in the double membrane system is in fact coupled. The correlated movement of the strand in the double membrane system is due to head-to-head interaction of claudins with those in a second membrane, which preserves the tetrameric structure of claudin pores.

**Figure 7:**
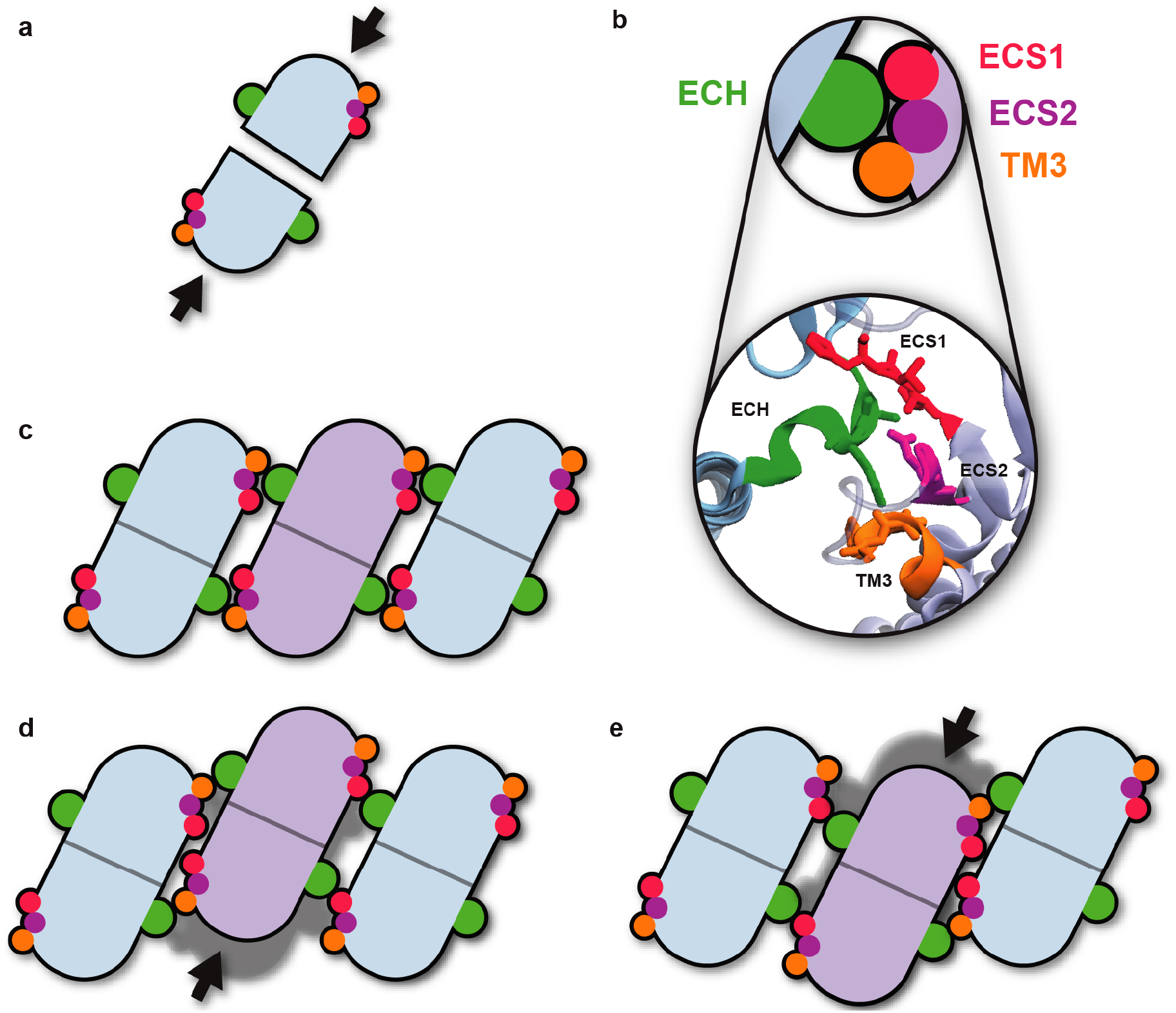
Flexibility mechanism of claudin-15 strands. Claudins monomers polymerize the cell membrane to form an antiparallel double row strand facilitated by (a) face-to-face *cis-*interactions and (b) side-by-side *cis-*interactions between monomers. The tetrameric ion channel pore is then formed by head-to-head (*trans-*) interactions between two claudin strands (not shown here). Three side-by-side interacting regions identified in the simulations are shown in (b). The extracellular helix ECH (green) acts as a pivot point as the local curvatures of the strand changes in the plane of membrane. (c-e) Claudin pairs connected through facte-to-face interactions slide with respect to each other to respond to local fluctuations in the strand curvature. Lateral flexibility of the strand originates from robustness of side-by-side *cis-*interactions via these three interacting regions.

## Conclusions

We have used molecular dynamics simulations to investigate mechanical flexibility of claudin-15 strands ranging from 30 to 225 nm in length. Claudin strands within tight junctions are dynamic and respond to external forces in epithelial cells by rearranging within the cell membrane and adapting their curvature. The paracellular barrier function is expected to remain intact during this process. We used our previously refined model of claudin-15 paracellular ion channels^40^ to simulate claudin strands with 36 to 300 monomers in two parallel lipid membranes. To present a less computationally yet accurate model of claudin strand assemblies we simulated the strands at all-atom as well as hybrid resolution, where the protein is represented atomically and lipid and water molecules are coarse-grained. Our results indicate that double strands of claudin-15 are dynamic in the plane of membrane with a persistence length of ∼174 nm consistent with experimental studies^12^

Our results reveal that functional flexibility of claudin-15 strands is due to the interplay of three sets of intermolecular interactions; the side-by-side *cis-*interactions between claudins in a row as observed in the crystal structure of claudin-15, the “face-to-face” interactions between *β*-sheets of claudins in two adjacent rows within the same membrane, and the head-to-head *trans-*interactions between extracellular domains of claudins in two membranes. The side-by-side interactions are the most flexible interaction network among the three. They provide rotational flexibility to neighboring claudins with a pivot point at the extracellular helix (ECH). The face-to-face and head-to-head interactions are least affected upon curving of the strands and are responsible for maintaining the tetrameric structure of the pore.

These finding suggest a novel mechanism for bending flexibility of claudins strands, in which claudins are assembled into interlocking tetrameric ion channels along the strand that slide with respect to each other as the strands curve over sub-micrometer length scales. The side-by-side interactions provide the structural flexibility for this movement while limiting its range. Water and ion density profiles indicate that during this process, the ion channels remain intact and paracellular seal is maintained. Thus, the barrier function is preserved.

These studies provide a mechanistic view on functional flexibility of tight junction strands and the role of claudin-claudin interactions in maintaining epithelial barrier function. Further studies are needed to understand large scale rearrangement of tight junction networks that are accompanied by formation of new tight junction branches, junctional strands, or involve other tight junction proteins in modulating strand flexibility.

## Materials and methods

### Molecular dynamics simulations

Molecular dynamics (MD) simulations were performed using the program NAMD^71^. All-atom simulations were carried out using CHARMM36 forcefield for proteins^72–74^, ions^75^, and phospholipids^76^ with the TIP3P water model^75^. These simulations were carried out with a time-step of 1 fs and assuming periodic boundary conditions. Langevin dynamics with a friction coefficient of *γ* = 5 ps^−1^ was used to keep temperature constant at 300 K. The Langevin Nosé-Hoover method^77^ was used to maintain the pressure at 1 atm in constant pressure simulations. Long-range electrostatic forces were calculated using the particle mech Ewald method^78^ with a grid spacing of at least 1 Å in each direction. The simulations used a time-step of 1 fs, 2 fs and 4 fs for bonded, short-range nonbonded and long-range electrostatic interactions calculations, respectively. A 1-4 rule is applied to the calculation of nonbonded interactions. The neighboring list was updated every 20 steps. The electrostatic interactions were switched to zero between 10 Å and 12 Å, and the van der Waals interactions cutoff is set to 12 Å. The distance cutoff for searching the lists of interacting pairs was set to 13.5 Å. Additional restraints were applied by enforcing a harmonic potential with a force constant of 2.0 kcal.mol^−1^.Å ^−2^, unless otherwise stated.

The hybrid resolution models were built and simulated using the PACE forcefield^44–46^. PACE forcefield models lipid and solvents consistent with the MARTINI forcefield^47^, while proteins are represented by a united-atom (UA) model, where heavy atoms and polar hydrogens are explicitly represented. The hybrid resolution models were simulated using a time-step of 2 fs or 3 fs, with the neighbor list being updated every 10 steps. The electrostatic interactions are switched to zero between 9 Å and 12 Å, and the van der Waals interactions cutoff is set to 12 Å. The distance cutoff for searching the lists of interacting pairs is set to 14 Å. Periodic boundary conditions were applied in all three directions. Langevin dynamics with a friction coefficient of *γ* = 5 ps^−1^ was used to keep the temperature constant at 300 K. The Langevin Nosé-Hoover method^77^ was used to maintain the pressure at 1 atm in constant pressure simulations. The dielectric constant is set to 15 as in Martini simulations with non-polarizable water^47^.

### System setup

Five models of claudin-15 strands in two parallel lipid bilayers were constructed. The initial structures of claudin-15 strands were based on our refined model of three claudin-15 pores (12 claudin-15 monomers) in two parallel POPC lipid bilayers^40^. In this model, each membrane contains six claudin-15 monomers that are arranged linearly into an antiparallel double row and form three half-pipe structures. Claudin pores are then formed by head-to-head interactions between extra-cellular domains of claudins in two parallel lipid bilayers, similar to the pores shown in Fig. 1. The three-pore claudin-15 model, equilibrated in POPC bilayers^40^, was replicated to build longer claudin strands consisting of 8, 20, 44 and 74 pores (each pore consists of four monomers) that were 27, 63, 135, and 225 nm long. Four patches of hydrated lipid bilayers (50 Å × 50 Å) were added on the two side of the claudin strand to separate the strand from its periodic image.

The all-atom models were then converted into a hybrid-resolution model based on the PACE forcefield^44–46^. The conversion is performed using the python scripts provided by Han’s laboratory at (http://web.pkusz.edu.cn/han/) to convert protein, lipids, and ions into the hybrid model. The system was then solvated by coarse-grained Martini water molecules^47^ using VMD^79^. The hybrid model systems consist of 200 K-1.1 M particles.

The smallest system, i.e., a system consisting of 36 claudin-15 monomers (8 pores) was simulated in all-atom representation, as well. This system was equilibrated for 300 ns in a multi-step process as described below. In the first step, the protein backbone and lipid head groups were restrained harmonically with a force constant of 2 kcal.mol^−1^.Å ^−2^ and the lipid tails were equilibrated for 200 ps at constant volume and temperature. In the next step, the lipid head groups were released and the system was equilibrated at constant pressure for 2 ns, after which, the two extracellular loops of claudins (residues 33–44 of ECS1 and residues 148–155 of ECS2) were released to move freely while the rest of the protein backbone was gradually released by decreasing the force constant to 1.0, 0.75, 0.5, and 0.25 kcal.mol^−1^.Å ^−2^ over 45 ns. After releasing all restraints, the system was equilibrated freely for 250 ns at constant pressure.

Four hybrid model systems containing 8, 20, 44, and 74 pores (36, 84, 180 and 300 claudin-15 monomers) were equilibrated for 1.125 *µ*s following a process similar to the equilibration of the allatom system. Briefly, by harmonically restraining the protein backbone and lipid head groups with a force constant of 2 kcal.mol^−1^.Å ^−2^, the lipid tails were equilibrated for 200 ps at constant volume and temperature. The lipid head groups were then released and the systems were equilibrated for 2 ns at constant pressure and temperature. Finally, the two extracellular loops of claudins (residues 33–44 of ECS1 and residues 148–155 of ECS2) were released to move freely while the rest of the protein backbone was gradually released by decreasing the force constant to 1.0, 0.75, 0.5, and 0.25 kcal.mol^−1^.Å ^−2^ over 70 ns. These equilibration processes were preformed at a time step of 2 fs. The simulations were then followed by 1.055 *µ*s of simulation time at constant pressure, using a time step of 2 fs for the first 140 ns and a time step of 3 fs for the last 915 ns.

### Persistence length calculations

By applying the worm-like chain approximation of linear polymers, persistence length (*l*_*p*_) of claudin-15 strands is directly calculated from thermal fluctuations of the strands in equilibrium MD simulation trajectories. To achieve this, the claudin strand is assumed to be a two dimensional linear polymer in the plane of membrane made of discrete segments. Each four pore-forming claudins are considered to be a segment located at position 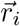, where 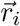, is the center of mass of the four claudins.

At each point on the strand 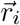, the tangent angle *ϕ*_*i*_ is determined as the angle corresponding to the tangent vector 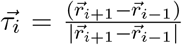. The persistence length is then calculated as the length of the strand over which the tangent angles become uncorrelated. Assuming a discrete strand made of segments with length *δs*, the contour length between any two points *i* and *j* on the strand is estimated to be *s* = |*i* − *j*|*δs*. We defined the contour fragment *δs* to be the distance between two segments 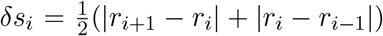 averaged over all segments during the simulation time; 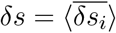.

The persistence length *l*_*p*_ is then calculated as the length over which the correlation between the tangent angles defined as ⟨cos(*ϕ*(*s*) − *ϕ*(0))⟩ is dropped by a factor of e:

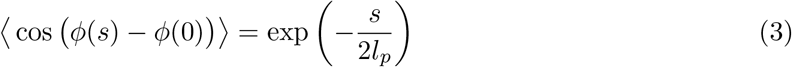

The factor of two in the denominator is due to the two dimensional nature of the strands. The persistence length is calculated for the five claudin-15 strand simulations as described above. The last 600 ns of the simulations trajectories are loaded at every 60 ps, and the center of mass of each four pore-forming claudins are recorded for each frame. The initial slopes of the decay of the logarithm of the cosine correlation functions are used to calculate the persistence length. The number of points included in curve fitting is determined by a R-squared cut-off of 0.95.

### Relative orientation and contact maps analyses

The relative orientation angle of adjacent claudins in the lipid membrane (*θ*) is calculated with respect to the initial arrangement of claudins in the crystal-structure (*θ* = 0). For each pair of neighboring claudins located at positions *i* and *i* + 1, a vector 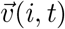 is defined connecting the center of mass of the backbone of residue 84 on TM2 of the first claudin (*i*) to the center of mass of the backbone of residue 170 on TM4 of the second claudin (*i* + 1) at time *t*. For each claudin *i*, the orientation angle *θ* is defined as the angle between 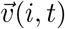 and 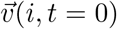, which corresponds to the crystallographic arrangement (Fig. 5). The last 600 ns of the simulation trajectory of the longest strand (225 nm with 300 claudins) was used for this analysis with a frequency of 60 ps, and the angles were reported for each of the 296 claudins after excluding the edges.

To characterize the side-by-side (*cis*-) interactions between adjacent claudins, a contact map of residues from neighboring claudins that are within 5.5 Å of each other is created at each frame and correlated with the relative orientation angles (*θ*). The last 15 ns of the simulation trajectory of the longest strand, i.e, the 225 nm strand (300 claudins) was used for this analysis. To be consistent with the other analyses, the trajectory was analyzed every 60 ps. Claudin pairs were clustered based on their relative orientation angle (*θ*) in three groups corresponding to −30° < *θ* < −10°, −10° < *θ* < 10°, and 10° < *θ* < 30°. Based on the residue-residue contact maps at each cluster, the probability of the dominant contacts were determined and the most dominant contacts are presented. The contacts with less than 20% probability of occurrence are not reported.

## Acknowledgements

The work of F.K.-A., S.F., S.M., and C.R.W. were supported by the National Science Foundation grant MCB-1846021. The authors acknowledge the Texas Advanced Computing Center (TACC) at The University of Texas at Austin for providing HPC resources that have contributed to the research results reported within this paper. URL: http://www.tacc.utexas.edu.

## References

1. V. W. Tang and D. A. Goodenough. Paracellular ion channel at the tight junction. Biophysical journal, 84:1660–1673, 2003.

2. J. M. Anderson and C. M. Van Itallie. Physiology and function of the tight junction. Cold Spring Harbor perspectives in biology, 1:a002584, 2009.

3. C. Zihni, C. Mills, K. Matter, and M. S. Balda. Tight junctions: from simple barriers to multifunctional molecular gates. Nature reviews Molecular cell biology, 17:564, 2016.

4. M. A. Odenwald and J. R. Turner. The intestinal epithelial barrier: a therapeutic target? Nature reviews Gastroenterology & hepatology, 14:9, 2017.

5. M. Furuse. Molecular basis of the core structure of tight junctions. Cold Spring Harbor perspectives in biology, 2:a002907, 2010.

6. L. A. Staehelin, T. Mukherjee, and A. W. Williams. Freeze-etch appearance of the tight junctions in the epithelium of small and large intestine of mice. Protoplasma, 67:165–184, 1969.

7. P. Claude and D. A. Goodenough. Fracture faces of zonulae occludentes from” tight” and” leaky” epithelia. The Journal of cell biology, 58:390–400, 1973.

8. S. Tsukita, M. Furuse, and M. Itoh. Multifunctional strands in tight junctions. Nature reviews Molecular cell biology, 2:285–293, 2001.

9. E. S. Krystofiak, J. B. Heymann, and B. Kachar. Carbon replicas reveal double stranded structure of tight junctions in phase-contrast electron microscopy. Communications biology, 2:1–6, 2019.

10. H. Sasaki, C. Matsui, K. Furuse, Y. Mimori-Kiyosue, M. Furuse, and S. Tsukita. Dynamic behavior of paired claudin strands within apposing plasma membranes. Proceedings of the National Academy of Sciences, 100:3971–3976, 2003.

11. C. M. Van Itallie, A. J. Tietgens, and J. M. Anderson. Visualizing the dynamic coupling of claudin strands to the actin cytoskeleton through zo-1. Molecular biology of the cell, 28:524–534, 2017.

12. J. Zhao, E. S. Krystofiak, A. Ballesteros, R. Cui, C. M. Van Itallie, J. M. Anderson, C. Fenollar-Ferrer, and B. Kachar. Multiple claudin–claudin cis interfaces are required for tight junction strand formation and inherent flexibility. Communications biology, 1:50, 2018.

13. A. M. Marchiando, L. Shen, W. V. Graham, K. L. Edelblum, C. A. Duckworth, Y. Guan, M. H. Montrose, J. R. Turner, and A. J. Watson. The epithelial barrier is maintained by in vivo tight junction expansion during pathologic intestinal epithelial shedding. Gastroenterology, 140:1208–1218, 2011.

14. T. Higashi, T. R. Arnold, R. E. Stephenson, K. M. Dinshaw, and A. L. Miller. Maintenance of the epithelial barrier and remodeling of cell-cell junctions during cytokinesis. Current biology, 26:1829–1842, 2016.

15. S. Varadarajan, R. E. Stephenson, and A. L. Miller. Multiscale dynamics of tight junction remodeling. Journal of cell science, 132, 2019.

16. P. Claude. Morphological factors influencing transepithelial permeability: A model for the resistance of thezonula occludens. The Journal of membrane biology, 39:219–232, 1978.

17. M. Furuse, K. Furuse, H. Sasaki, and S. Tsukita. Conversion of zonulae occludentes from tight to leaky strand type by introducing claudin-2 into madin-darby canine kidney i cells. The Journal of cell biology, 153:263–272, 2001.

18. C. R. Weber, D. R. Raleigh, L. Su, L. Shen, E. A. Sullivan, Y. Wang, and J. R. Turner. Epithelial myosin light chain kinase activation induces mucosal interleukin-13 expression to alter tight junction ion selectivity. Journal of Biological Chemistry, 285:12037–12046, 2010.

19. M. Furuse, T. Hirase, M. Itoh, A. Nagafuchi, S. Yonemura, S. Tsukita, and S. Tsukita. Occludin: a novel integral membrane protein localizing at tight junctions. The Journal of cell biology, 123:1777–1788, 1993.

20. Y. Ando-Akatsuka, M. Saitou, T. Hirase, M. Kishi, A. Sakakibara, M. Itoh, S. Yonemura, M. Furuse, and S. Tsukita. Interspecies diversity of the occludin sequence: cdna cloning of human, mouse, dog, and rat-kangaroo homologues. The Journal of cell biology, 133:43–47, 1996.

21. Martìn-Padura, S. Lostaglio, M. Schneemann, L. Williams, M. Romano, P. Fruscella, C. Panzeri, A. Stoppacciaro, L. Ruco, A. Villa, et al. Junctional adhesion molecule, a novel member of the immunoglobulin superfamily that distributes at intercellular junctions and modulates monocyte transmigration. The Journal of cell biology, 142:117–127, 1998.

22. M. Furuse, K. Fujita, T. Hiiragi, K. Fujimoto, and S. Tsukita. Claudin-1 and -2: novel integral membrane proteins localizing at tight junctions with no sequence similarity to occludin. J. Cell Biol, 141:1539–1550, 1998.

23. M. Furuse, H. Sasaki, K. Fujimoto, and S. Tsukita. A single gene product, claudin-1 or-2, reconstitutes tight junction strands and recruits occludin in fibroblasts. The Journal of cell biology, 143:391–401, 1998.

24. M. Furuse and S. Tsukita. Manner of interaction of heterogeneous cluadin species within and between tight junction strands. J. Cell. Biol., 147:891–903, 1999.

25. C. M. Van Itallie and J. M. Anderson. Claudins and epithelial paracellular transport. Annu. Rev. Physiol., 68:403–429, 2006.

26. S. Angelow, R. Ahlstrom, and A. S. Yu. Biology of claudins. American Journal of Physiology-Renal Physiology, 295:F867–F876, 2008.

27. D. Gunzel and A. S. L. Yu. Claudins and the modulation of tight junction permeability. Physiol. Rev., 93:525–569, 2013.

28. S. Tsukita, H. Tanaka, and A. Tamura. The claudins: from tight junctions to biological systems. Trends in biochemical sciences, 44:141–152, 2019.

29. Piontek, S. M. Krug, J. Protze, G. Krause, and M. Fromm. Molecular architecture and assembly of the tight junction backbone. Biochimica et Biophysica Acta (BBA)-Biomembranes, 1862:183279, 2020.

30. O. R. Colegio, C. M. Van Itallie, H. J. McCrea, C. Rahner, and J. M. Anderson. Claudins create charge-selective channels in the paracellular pathway between epithelial cells. Am. J. Physiol. Cell Physiol., 283:C142–C147, 2002.

31. A. S. Yu, M. H. Cheng, S. Angelow, D. Günzel, S. A. Kanzawa, E. E. Schneeberger, M. Fromm, and R. D. Coalson. Molecular basis for cation selectivity in claudin-2–based paracellular pores: identification of an electrostatic interaction site. The Journal of general physiology, 133:111–127, 2009.

32. H. Suzuki, T. Nishizawa, K. Tani, Y. Yamazaki, A. Tamura, R. Ishitani, N. Dohmae, S. Tsukita, O. Nureki, and Y. Fujiyoshi. Crystal structure of a claudin provides insight into the architecture of tight junctions. Science, 344:304–307, 2014.

33. Li, S. Angelow, A. Linge, M. Zhuo, and A. S. Yu. Claudin-2 pore function requires an intramolecular disulfide bond between two conserved extracellular cysteines. American Journal of Physiology-Cell Physiology, 305:C190–C196, 2013.

34. H. Suzuki, K. Tani, A. Tamura, S. Tsukita, and Y. Fujiyoshi. Model for the architecture of claudin-based paracellular ion channels through tight junctions. Journal of molecular biology, 427:291–297, 2015.

35. G. Krause, J. Protze, and J. Piontek. Assembly and function of claudins: Structure–function relationships based on homology models and crystal structures. In Seminars in cell & developmental biology, volume 42, pages 3–12. Elsevier, 2015.

36. J. Piontek, L. Winkler, H. Wolburg, S. Muller, N. Zuleger, C. Piehl, B. Wiesner, G. Krause, and I. Blasig. Formation of tight junction: Determinants of hemophilic interaction between classic claudins. FASEB J., 22:146–158, 2008.

37. A. Piontek, J. Rossa, J. Protze, H. Wolburg, C. Hempel, D. Günzel, G. Krause, and J. Piontek. Polar and charged extracellular residues conserved among barrier-forming claudins contribute to tight junction strand formation. Annals of the New York Academy of Sciences, 1397:143–156, 2017.

38. S. Angelow and A. S. Yu. Structure-function studies of claudin extracellular domains by cysteine-scanning mutagenesis. Journal of Biological Chemistry, 284:29205–29217, 2009.

39. J. Rossa, C. Ploeger, F. Vorreiter, T. Saleh, J. Protze, D. Günzel, H. Wolburg, G. Krause, and J. Piontek. Claudin-3 and claudin-5 protein folding and assembly into the tight junction are controlled by non-conserved residues in the transmembrane 3 (tm3) and extracellular loop 2 (ecl2) segments. Journal of Biological Chemistry, 289:7641–7653, 2014.

40. P. Samanta, Y. Wang, S. Fuladi, J. Zou, Y. Li, L. Shen, C. Weber, and F. Khalili-Araghi. Molecular determination of claudin-15 organization and channel selectivity. The Journal of general physiology, 150:949–968, 2018.

41. G. Alberini, F. Benfenati, and L. Maragniano. A refined model of claudin-15 tight junction paracellular architecture by molecular dynamics simulations. PLoS One, 12:e0184190, 2017.

42. G. Alberini, F. Benfenati, and L. Maragniano. Molecular dynamics simulations of ion selectivity in a claudin-15 paracellular channel. Journal of Physical Chemistry B, 122:10783–10792, 2018.

43. S. Fuladi, R.-W. Jannat, L. Shen, C. R. Weber, and F. Khalili-Araghi. Computational modeling of claudin structure and function. International journal of molecular sciences, 21:742, 2020.

44. W. Han, C.-K. Wan, F. Jiang, and Y.-D. Wu. Pace force field for protein simulations. 1. full parameterization of version 1 and verification. Journal of chemical theory and computation, 6:3373–3389, 2010.

45. C.-K. Wan, W. Han, and Y.-D. Wu. Parameterization of pace force field for membrane environment and simulation of helical peptides and helix–helix association. Journal of chemical theory and computation, 8:300–313, 2011.

46. W. Han and K. Schulten. Further optimization of a hybrid united-atom and coarse-grained force field for folding simulations: improved backbone hydration and interactions between charged side chains. Journal of chemical theory and computation, 8:4413–4424, 2012.

47. S. J. Marrink, H. J. Risselada, S. Yefimov, D. P. Tieleman, and A. H. De Vries. The martini force field: coarse grained model for biomolecular simulations. The journal of physical chemistry B, 111:7812–7824, 2007.

48. T. S. Lim, R. S. V. dn W. Hunziker, and C. T. Lim. Kinetics of adhesion mediated by extracelluular loops of claudin-2 as revealed by single-molecule force spectroscopy. J. Mol. Biol., 381:681–691, 2008.

49. T. S. Lim, S. R. K. Vedula, S. Hui, P. J. Kausalya, W. Hunziker, and C. T. Lim. Probing effects of ph change on dynamic response of claudin-2 mediated adhesion using single molecule force spectroscopy. Experimental cell research, 314:2643–2651, 2008.

50. C. Piehl, J. Piontek, J. Cording, H. Wolburg, and I. Blasig. Participation of the second extracellular loop of claudin-5 in paracellular tightening against ions, small and large molecules. Cell Mol Life Sci, 67:2131–2140, 2010.

51. C. Frontali, E. Dore, A. Ferrauto, E. Gratton, A. Bettini, M. Pozzan, and E. Valdevit. An absolute method for the determination of the persistence length of native dna from electron micrographs. Biopolymers: Original Research on Biomolecules, 18:1353–1373, 1979.

52. N. Borochov, H. Eisenberg, and Z. Kam. Dependence of dna conformation on the concentration of salt. Biopolymers: Original Research on Biomolecules, 20:231–235, 1981.

53. G. S. Manning. The persistence length of dna is reached from the persistence length of its null isomer through an internal electrostatic stretching force. Biophysical journal, 91:3607–3616, 2006.

54. S. Geggier, A. Kotlyar, and A. Vologodskii. Temperature dependence of dna persistence length. Nucleic acids research, 39:1419–1426, 2011.

55. S. Brinkers, H. R. Dietrich, F. H. de Groote, I. T. Young, and B. Rieger. The persistence length of double stranded dna determined using dark field tethered particle motion. The Journal of chemical physics, 130:06B607, 2009.

56. S. B. Smith and A. J. Bendich. Electrophoretic charge density and persistence length of dna as measured by fluorescence microscopy. Biopolymers: Original Research on Biomolecules, 29:1167–1173, 1990.

57. F. Gittes, B. Mickey, J. Nettleton, and J. Howard. Flexural rigidity of microtubules and actin filaments measured from thermal fluctuations in shape. The Journal of cell biology, 120:923–934, 1993.

58. P. Venier, A. C. Maggs, M.-F. Carlier, and D. Pantaloni. Analysis of microtubule rigidity using hydrodynamic flow and thermal fluctuations. Journal of biological chemistry, 269:13353–13360, 1994.

59. D. Sept and F. C. MacKintosh. Microtubule elasticity: connecting all-atom simulations with continuum mechanics. Physical review letters, 104:018101, 2010.

60. Kikumoto, M. Kurachi, V. Tosa, and H. Tashiro. Flexural rigidity of individual microtubules measured by a buckling force with optical traps. Biophysical journal, 90:1687–1696, 2006.

61. H. Felgner, R. Frank, and M. Schliwa. Flexural rigidity of microtubules measured with the use of optical tweezers. Journal of cell science, 109:509–516, 1996.

62. J.-W. Chu and G. A. Voth. Allostery of actin filaments: molecular dynamics simulations and coarse-grained analysis. Proceedings of the National Academy of Sciences, 102:13111–13116, 2005.

63. A. Ott, M. Magnasco, A. Simon, and A. Libchaber. Measurement of the persistence length of polymerized actin using fluorescence microscopy. Physical Review E, 48:R1642, 1993.

64. C. M. Van Itallie, A. J. Tietgens, and J. M. Anderson. Visualizing the dynamic coupling of claudin strands to the actin cytoskeleton through zo-1. Molecular biology of the cell, 28:524–534, 2017.

65. R. M. Venable, F. L. Brown, and R. W. Pastor. Mechanical properties of lipid bilayers from molecular dynamics simulation. Chemistry and physics of lipids, 192:60–74, 2015.

66. P. W. Fowler, J. Hélie, A. Duncan, M. Chavent, H. Koldsø, and M. S. Sansom. Membrane stiffness is modified by integral membrane proteins. Soft Matter, 12:7792–7803, 2016.

67. D. Bochicchio and L. Monticelli. The membrane bending modulus in experiments and simulations: a puzzling picture. Advances in Biomembranes and Lipid Self-Assembly, 23:117–143, 2016.

68. A. F. Loftus, S. Noreng, V. L. Hsieh, and R. Parthasarathy. Robust measurement of membrane bending moduli using light sheet fluorescence imaging of vesicle fluctuations. Langmuir, 29:14588–14594, 2013.

69. S. Jablin, K. Akabori, and J. Nagle. Experimental support for tilt-dependent theory of biomembrane mechanics. Physical review letters, 113:248102, 2014.

70. G. Krause, L. Winkler, S. L. Mueller, R. F. Haseloff, J. Piontek, and I. E. Blasig. Structure and function of claudins. Biochim. Biophys. Acta, 1778:631–645, 2008.

71. J. Phillips, R. Braun, W. Wang, J. Gumbart, E. Tajkhorshid, E. Villa, C. Chipot, R. Skeel, L. Kale, and K. Schulten. Scalable molecular dynamics with NAMD. J. Comput. Chem., 26:1781–1802, 2005.

72. A. J. MacKerell, D. Bashford, M. Bellot, R. L. Dunbrack, J. Evanseck, M. Field, S. Fischer, J. Gao, H. Guo, D. Joseph-McCarthy, S. Ha, L. Kuchnir, K. Kuczera, F. Lau, C. Mattos, S. Michnick, T. Ngo, D. Nguyen, B. Prodhom, W. E. R. Iii, B. Roux, M. Schlenkrich, J. Smith, R. Stote, J. Straub, M. Watanabe, J. Wiorkiewicz-Kuzcera, and M. Karplus. All-atom empirical potential for molecular modeling and dynamics studies of proteins. J. Phys. Chem. B, 102:3586–3616, 1998.

73. A. D. Mackerell Jr, M. Feig, and C. L. Brooks III. Extending the treatment of backbone energetics in protein force fields: limitations of gas-phase quantum mechanics in reproducing protein conformational distributions in molecular dynamics simulations. Journal of computational chemistry, 25:1400–1415, 2004.

74. R. B. Best, X. Zhu, J. Shim, P. E. Lopes, J. Mittal, M. Feig, and A. D. MacKerell Jr. Optimization of the additive charmm all-atom protein force field targeting improved sampling of the backbone ϕ, ψ and side-chain χ1 and χ2 dihedral angles. Journal of chemical theory and computation, 8:3257–3273, 2012.

75. W. L. Jorgensen, J. Chandrasekhar, J. D. Madura, R. W. Impey, and M. L. Klein. Comparison of simple potential functions for simulating liquid water. The Journal of chemical physics, 79:926–935, 1983.

76. J. B. Klauda, R. M. Venable, J. A. Freites, J. W. O?Connor, D. J. Tobias, C. Mondragon-Ramirez, I. Vorobyov, A. D. MacKerell Jr, and R. W. Pastor. Update of the charmm all-atom additive force field for lipids: validation on six lipid types. The journal of physical chemistry B, 114:7830–7843, 2010.

77. S. E. Feller, Y. Zhang, R. W. Pastor, and B. R. Brooks. Constant pressure molecular dynamics simulation: the langevin piston method. The Journal of chemical physics, 103:4613–4621, 1995.

78. T. Darden, D. York, and L. Pedersen. Particle mesh ewald: An N.log(N) method for Ewald sums in large systems. J. Chem. Phys., 98:10089, 1993.

79. W. Humphrey, A. Dalke, and K. Schulten. Vmd: visual molecular dynamics. J. Mol. Graphics, 14:33–38, 1996.

