## Supplemental Information for "Molecular mechanism of claudin-15 strand flexibility"

Table 1: **Structural stability and RMSD values of claudin-15 strands.** Average RMSD of claudin-15 monomers are measured over 800 ns for hybrid-resolution strands and over 165 ns for the all-atom strand model. The errors are reported as the standard deviation.

| Claudin-15 Strands | | protein backbone RMSD (Å) | TM helices backbone RMSD (Å) |
|---|---|---|---|
| Hybrid-resolution model | 300 cldn – 225 nm | 3.92 ± 0.85 | 1.61 ± 0.55 |
| | 180 cldn – 135 nm | 3.99 ± 0.94 | 1.66 ± 0.63 |
| | 84 cldn – 63 nm | 4.05 ± 1.01 | 1.66 ± 0.63 |
| | 36 cldn – 27 nm | 4.21 ± 0.88 | 1.76 ± 0.53 |
| All-atom model | 36 cldn – 27 nm | 2.19 ± 0.49 | 0.81 ± 0.16 |

Table 2: **Structural stability and RMSD values of claudin-15 pores.** Average RMSD of claudin-15 pores formed by 8 interacting monomers are measured over 800 ns for hybrid-resolution models and over 165 ns for the all-atom strand model. The errors are reported as the standard deviation.

| Claudin-15 Pores | | Protein backbone RMSD (Å) | TM helices backbone RMSD (Å) |
|---|---|---|---|
| Hybrid-resolution model | 74 pores – 225 nm | 5.97 ± 0.79 | 5.18 ± 0.93 |
| | 44 pores – 135 nm | 6.14 ± 0.86 | 5.33 ± 0.95 |
| | 20 pores – 63 nm | 6.53 ± 0.92 | 5.75 ± 0.97 |
| | 8 pores – 27 nm | 6.56 ± 0.89 | 5.79 ± 1.15 |
| All-atom model | 8 pores – 27 nm | 3.86 ± 0.30 | 3.39 ± 0.33 |
